# Linkage mapping and genome annotation give novel insights into gene family expansions and regional recombination rate variation in the painted lady (*Vanessa cardui*) butterfly

**DOI:** 10.1101/2022.04.14.488360

**Authors:** Daria Shipilina, Karin Näsvall, Lars Höök, Roger Vila, Gerard Talavera, Niclas Backström

## Abstract

Gene family expansions and crossing over are two main mechanisms for the generation of novel genetic variants that can be picked up by natural selection. Here, we developed a high-density, pedigree-based linkage map of the painted lady butterfly (*Vanessa cardui*) – a non-diapausing, highly polyphagous species famous for its long-distance migratory behavior. We also performed detailed annotations of genes and interspersed repetitive elements for a previously developed genome assembly, characterized species-specific gene family expansions and the relationship between recombination rate variation and genomic features. Identified expanded gene families consisted of clusters of tandem duplications with functions associated with protein and fat metabolism, detoxification, and defense against infection - key functions for the painted lady’s unique lifestyle. The detailed assessment of recombination rate variation demonstrated a negative association between recombination rate and chromosome size. Moreover, the recombination landscape along the holocentric chromosomes was bimodal. The regional recombination rate was positively associated with the proportion of short interspersed elements (SINEs), but not the other repeat classes, potentially a consequence of SINEs hijacking the recombination machinery for proliferation. The detailed genetic map developed here will contribute to the understanding of the mechanisms and evolutionary consequences of recombination rate variation in Lepidoptera in general. We conclude that the structure of the painted lady genome has been shaped by a complex interplay between recombination, gene duplications and TE-activity and that specific gene family expansions have been key for the evolution of long-distance migration and the ability to utilize a wide range of host plants.

## Introduction

The genomic era opens up opportunities for investigating relationships between genotypes and complex phenotypes on a novel level and for a better understanding of genome evolution. Combinations of different approaches can lead to novel insights into the dynamics of recurring duplications, deletions and other types of structural rearrangements, for example, by assessing molecular mechanisms and evolutionary consequences of gene family expansions and contractions, the activity of selfish genetic elements (e.g. transposable elements, TEs) and recombination rate variation.

Gene duplication has since long been recognised as an important mechanism for generating novel genetic material for natural selection to act upon (Henikoff, 1997; Ojeda-López et al., 2020; Zhang, 2003), and gene family expansions and contractions are important sources for generation of phenotypic diversity (Chen et al., 2013; Kondrashov, 2012). Comparative approaches, such as orthology analysis, allow for identification of expanding or contracting gene families and annotation of specific orthogroups (i.e. gene sets originating from a single gene copy in the common ancestor of the focal taxa) can aid in the assessment of the functional relevance of gene copy number variation in the evolution of lineage-specific traits. This approach might be beneficial for investigating complex phenotypes, where combined effects of different types of genetic changes likely underlie the trait (Schwander et al., 2014). Analysis of gene expansion dynamics can complement traditional studies of sequence changes in single orthologous genes, especially when comparing distant taxa.

Since the spearheading work by McClintock (McClintock, 1956), transposable elements (TEs) have been acknowledged as major contributors to different types of evolutionary change in eukaryotes (Kazazian, 2004; Kidwell and Lisch, 1997). Transposable elements are capable of self-replication within the host genome, resulting in the presence of multiple interspersed copies, which in turn can mediate both small scale deletions and duplications and large scale chromosome rearrangements (Kidwell and Lisch, 1997). In addition, TE insertions can affect gene function when regulatory or coding regions are targeted. Although this can sometimes lead to an instant selective advantage (Van’t Hof et al., 2016), TE propagations predominantly have neutral or deleterious effects on the host (Hedges and Deininger, 2007). The proliferation of TEs in the genome of a host organism can be considered a selfish process since TEs enhance their own transmission, a process that leads to the rapid diversification of many eukaryotic genomes (Wells and Feschotte, 2020). In *Heliconius* butterflies, TE diversification has likely contributed to the formation of reproductive barriers (Ray et al., 2019), and in other butterfly lineages, rapid accumulation of TEs has resulted in considerable genome size expansions (Podsiadlowski et al., 2021; Talla et al., 2017). Therefore, characterisation of the TE repertoire is key to understanding the microevolutionary dynamics within the genome of a species and the potential effects of TE activity on trait variation within populations and between species.

Besides gene duplication and TE activity, recombination is a process crucial for evolutionary innovation. Meiotic recombination shuffles existing segregating genetic variants, resulting in the generation of novel haplotypes (Peñalba and Wolf, 2020). Recombination also influences selection efficiency by directly preventing the accumulation of deleterious alleles (Müller’s ratchet) and breaking the physical linkage between mutations with different selective effects (Hill-Robertson effects). The rate of recombination can vary on different scales. Of particular interest for population genetic processes is the variation in recombination rate between different genomic regions. Such spatial variation in the recombination rate has been observed in many different organisms (Stapley et al., 2017; Tiley and Burleigh, 2015). However, besides detailed recombination maps in the butterfly genus *Heliconius* (Martin et al., 2019), little is known about how the rate of recombination rate varies across chromosome regions in Lepidoptera and how recombination is associated with different genomic features (Haenel et al., 2018; Talla et al., 2019).

As indicated above, incorporating different approaches is essential for studying the genetic underpinnings of complex phenotypes and the mechanisms governing microevolutionary processes. The painted lady, *Vanessa cardui*, represents a key study system for a wide array of evolutionary studies. It is the most wide-spread of all butterfly species (Talavera et al., 2018), and its migratory behavior includes a diverse repertoire of distinct phenotypes. In general, migratory butterfly species need to sustain long-distance flight and have well developed navigational abilities (Chapman et al., 2015; Guerra et al., 2014). Therefore, traits related to energy metabolism, sensory reception and the flight machinery have likely been under strong directional selection. In contrast to many other migratory butterflies inhabiting temperate zones, the painted lady is a non-diapausing, multigenerational migrant, with an annual migratory circuit covering areas with extreme environmental heterogeneity (Menchetti et al., 2019; Talavera and Vila, n.d.). Despite the high risks associated with such a migratory lifestyle, painted ladies have successfully colonized almost all continents, and the species harbors high levels of genetic diversity, indicating a large effective population size (Garcìa-Berro et al., in prep). This could also be a consequence of the species’ ability to utilize a wide range of host plants (Ackery, 1988; Celorio-Mancera et al., n.d.). Until the era of high-throughput sequencing, the possibilities to gain insights into how the migratory and generalist lifestyle has been manifested at the level of the genome have been limited: genetic basis of migratory behavior in insects has only been investigated in a few model species (e.g. the monarch butterfly, *Danaus plexippus*) so far (Merlin and Liedvogel, 2019).

A key step for genomic analyses is the development of a high-contiguity genome assembly of the focal species and a thorough genome annotation. A powerful method to ensure the spatial correctness of a chromosome level physical assembly is construction of a linkage map. In this study, we present the first detailed linkage map of the painted lady and verify scaffolds from a previously available genome assembly based on long-read sequencing technology (Lohse et al., 2021). We use the genome annotation and linkage information to quantify lineage-specific patterns of gene family evolution, relative TE abundance and how the regional recombination rate variation is associated with genomic features in the painted lady. Our analyses complement earlier efforts to establish genomic tools for this species (Connahs et al., 2016; Zhang et al., 2021) and give novel insights into the overall genome structure, recombination rate variation and lineage-specific gene family expansions in this species, information that informs on the molecular mechanisms underlying genome evolution in butterflies in general and the formation of the complex migratory phenotype and generalist lifestyle of the painted lady in particular.

## Results

### Linkage map and genome annotation

To verify a chromosome level assembly of the painted lady (Lohse et al., 2021) and to get access to detailed recombination rate data, we constructed a pedigree-based linkage map. The total distance of the linkage map was 1,516 centiMorgan (cM) and contained 1,323 markers. When anchored on the 424 Mb physical assembly, the average marker density was 3.09 markers / Mb. The genome assembly was highly collinear with the marker order in the linkage map (Pearson’s correlation coefficient; R = 0.91-1.00, p-value > 1.00×10^−04^, Figure S1) and consisted of 30 autosomes and the sex chromosomes Z and W. The high collinearity between linkage groups and assembled scaffolds, the large scaffold N50 (14.6 Mb) and high BUSCO scores (97% complete arthropod genes) confirm that the scaffolds in the assembly essentially represent complete chromosomes that could be used for accurate characterization of genomic features and quantification of regional recombination rate estimates.

In total, TEs constituted > 150 Mb (37.40%) of the assembly and LINEs and SINE were the most abundant of the characterized repeat classes (Table 1). After automatic annotation and subsequent manual curation, 13,161 protein-coding genes were identified (including 89.90% BUSCO genes), of which 12,209 had functional annotation information (Table 1). Visual inspection of the spatial distribution of genes and Tes along chromosomes revealed rather similar distributions of repeat classes between autosomes and the Z-chromosome, but also an observable excess of repeats on smaller autosomes and a striking difference in repeat composition and gene density on the W-chromosome (Figure 1).

**Table 1.** Linkage map, genome assembly and annotation statistics.

| Linkage map |  |
| --- | --- |
| Total map length (cM) | 1,516 |
| Number of markers | 1,323 |
| Markers per physical distance (N / Mb) | 3.09 |
| Genome assembly |  |
| Scaffold N50 (bp) | 14,615,999 |
| GC content | 33.41% |
| Total repeat proportion | 37.40% |
| Repeat content (% of total repeat proportion) |  |
| SINEs | 7.30% |
| LINEs | 14.94% |
| LTR elements | 2.47% |
| DNA elements | 3.04% |
| Simple and unknown repeats | 30.17% |
| Gene annotation |  |
| BUSCO genes | 89.90% |
| Number of protein coding genes | 13,161 |
| Number of genes with functional annotation | 12,209 |

**Figure 1.**
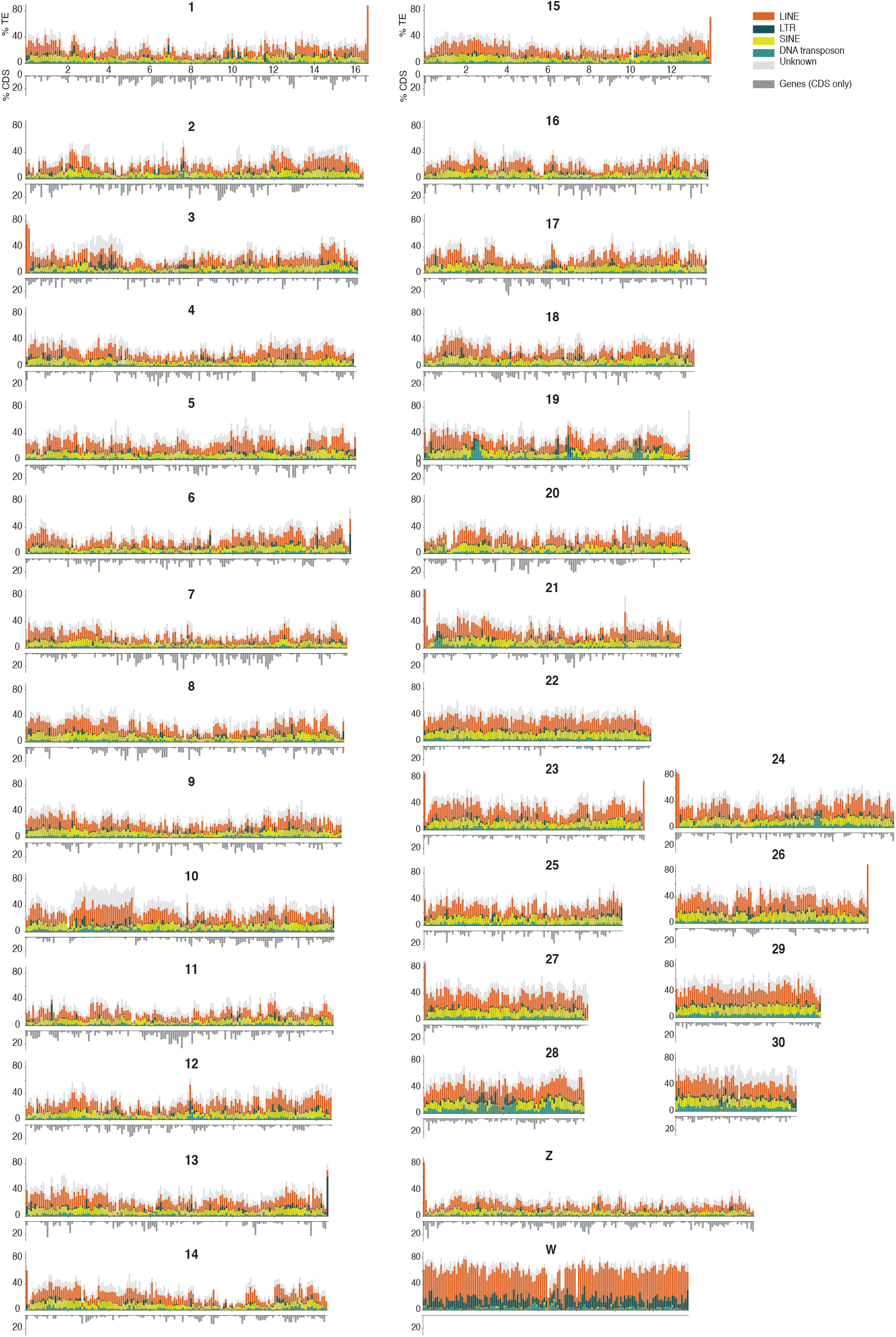
Distribution of repeat classes and genes as estimated along the painted lady chromosomes (100 kb windows). Density (% of the window covered) of different TE classes are illustrated with distinct colors cumulatively added on top of each other above the X-axis and density of genes below the X-axis (legend to the top right).

### Synteny

The level of large-scale structural conservation of the painted lady genome was assessed by comparing gene order on the painted lady chromosomes to two previously available high-contiguity lepidopteran genome assemblies positioned at different levels of divergence in the lepidopteran tree of life, the silkmoth (*Bombyx mori*) and the postman butterfly (*Heliconius melpomene*). Overall, the synteny was highly conserved between the painted lady and the other species, but chromosomes 28 and 26 mapped to the same chromosome (24) in *B. mori* and the previously described fusions of several chromosomes in the *H. melpomene* genome (Davey et al., 2017) could also be verified (Figure 2). In summary, this confirms that the painted lady karyotype is highly similar to the inferred ancestral butterfly karyotype (Ahola et al., 2014).

**Figure 2.**
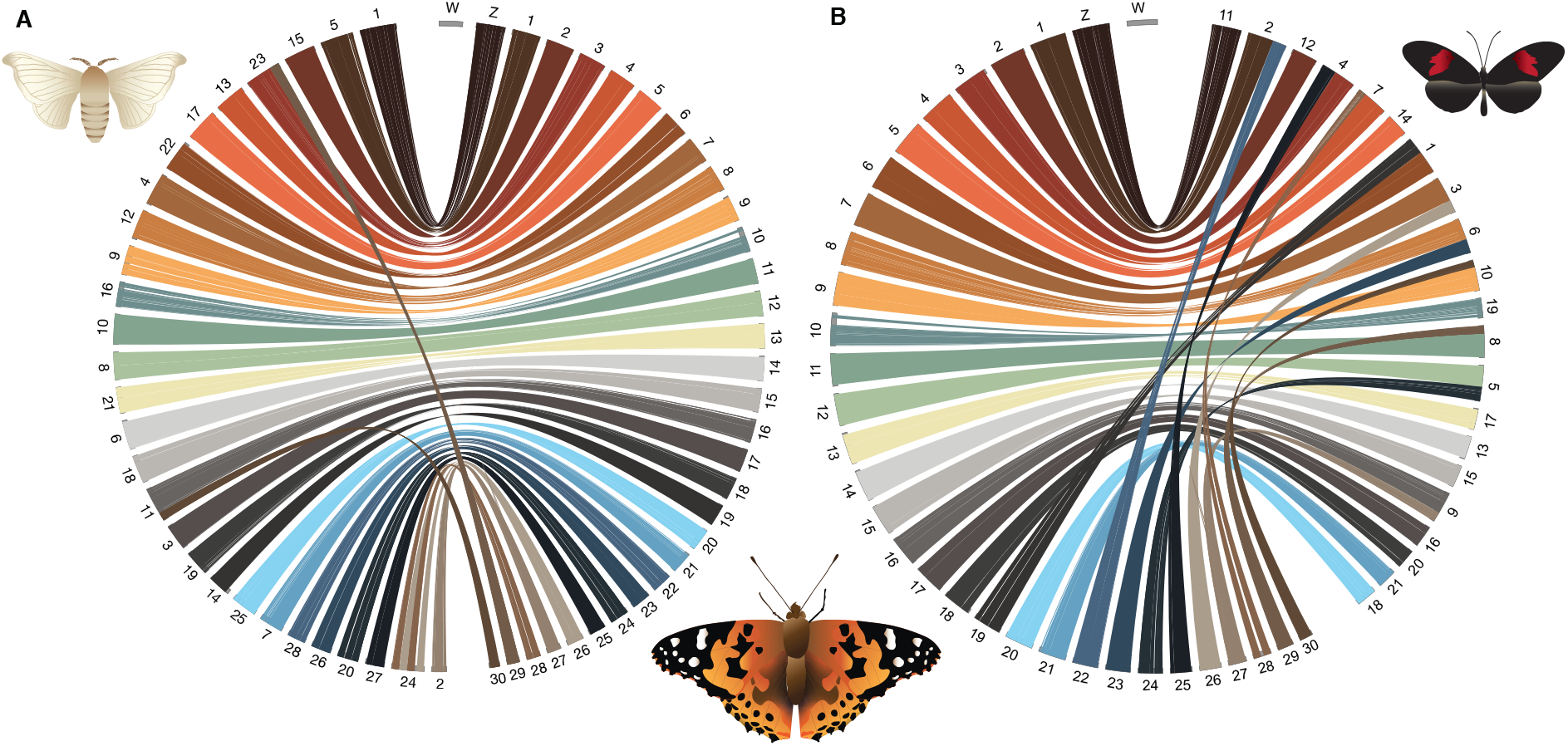
Synteny between the painted lady and a) the silkmoth (*Bombyx mori*) and b) the postman butterfly (*Heliconius melpomene*) chromosomes, respectively. The painted lady autosomes are sorted and named according to length and the sex chromosomes are indicated with Z and W. Orthologous genes are connected with lines. Colors represent individual chromosomes.

### Gene family evolution

To investigate the turnover of specific gene families in the painted lady, we analyzed a set of nine representative nymphalid species with detailed annotation information (see methods). The non-migratory Kamehameha butterfly (*Vanessa tameamea*) was included to assess differences in gene family evolution between sedentary and migratory lineages within the *Vanessa* genus. We found that 93.2% (1,288,332) of the total number of genes from the nine nymphalid species were clustered in 14,027 orthogroups. The percentage of genes assigned to orthogroups varied from 86.7 to 99.6% in the different species (Table S1). In the painted lady, 96.4% (12,692) of the annotated genes were assigned to 10,361 orthogroups with 19 lineage-specific orthogroups containing 63 genes (Table S1). Within the *Vanessa* genus, 65 expansions had occurred on the ancestral branch, 648 on the *V. cardui* branch and 1,563 on the *V. tameamea* branch.

We used a maximum likelihood model to detect genes with distinct gene family expansion rates in the painted lady compared to the other species. The analysis showed that 15 orthogroups were significantly expanded in the painted lady. These orthogroups contained 77 genes, of which 34 had associated GO-terms (Figure 3). Among the largest expanded gene families were two classes of proteases, a lipoprotein receptor and the Lepidoptera-specific moricin immune-gene family. Analysis of the spatial distribution of extended orthogroups revealed clustering/tandem duplications for all except one of the orthogroups (Figure 3C). Significantly enriched GO-terms for expanded gene families in the painted lady were predominantly associated with protein degradation, muscle function and development, and fatty acid and energy metabolism (Figure 3A). Multiple ontology terms were shared between expanded orthogroups, pointing towards similar functions associated with the different gene families (Figure 3B). Additionally, we identified gene families with a distinct gene expansion rate in both the painted lady and the monarch butterfly *Danaus plexippus* - the latter a key model organism for insect migration studies - and compared those to the other nymphalids. This analysis revealed 11 orthogroups with a higher expansion rate and 29 orthogroups with genes specific to these two lineages. The common orthogroups included 112 genes and were significantly enriched for GO-terms predominantly associated with metabolic processes, defence against infection and neuronal activity (Figure S2).

**Figure 3.**
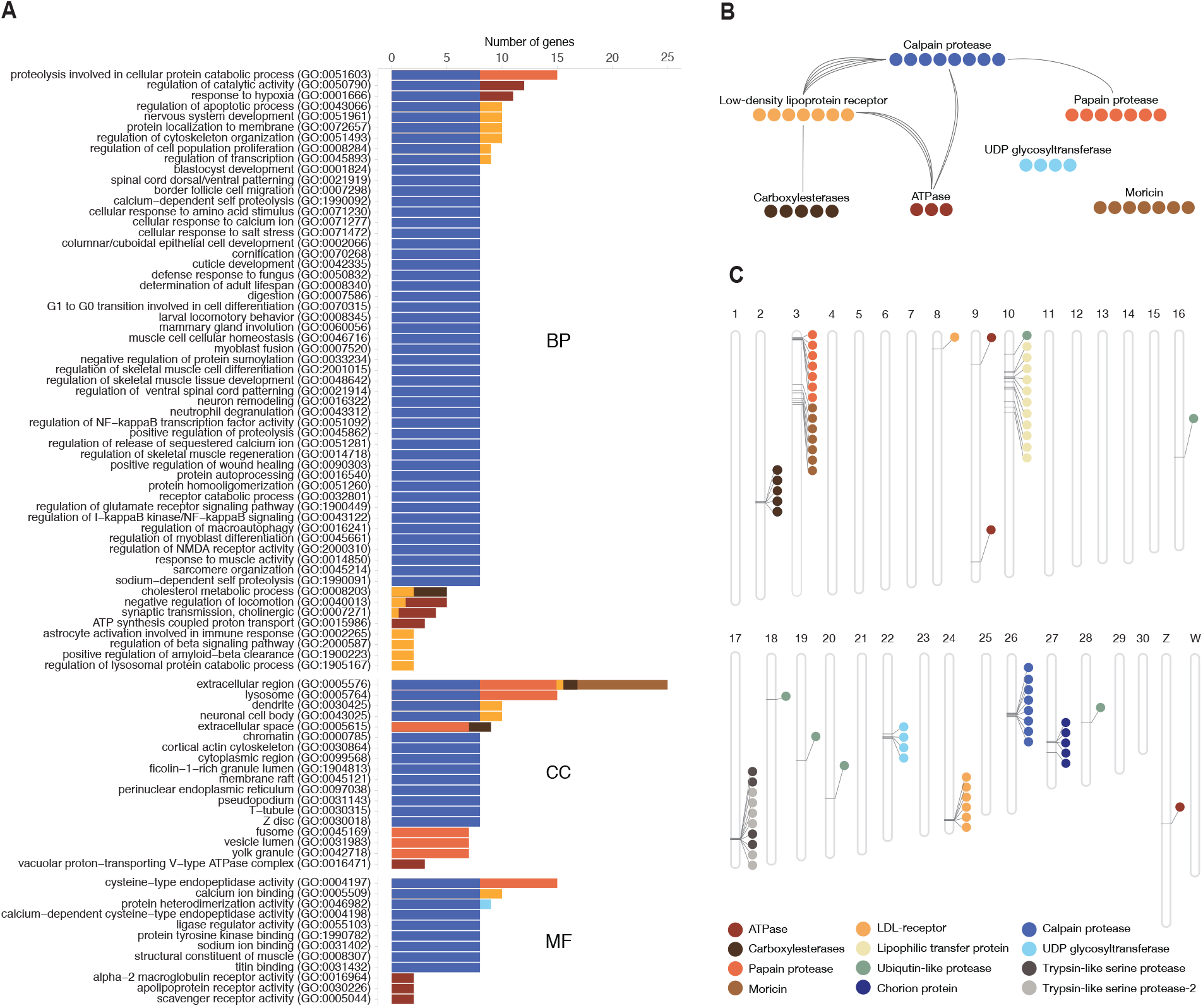
A) Significantly (p-value *<* 0.05 after FDR-correction) enriched gene ontology (GO) terms associated with expanded gene families in the painted lady. The bars show the number of genes associated with each GO-term. The different GO-categories are biological process (BP), cellular compartment (CC) and molecular function (MF). B) Orthogroups with significantly enriched GO-terms. Shared GO terms between ontology terms (biological process category only) are shown in connecting lines. C) Spatial distribution of genes from extended orthogroups identified in BadiRate analysis.

### Patterns of recombination rate variation

#### Global and chromosome specific recombination rates

The development of a detailed linkage map allowed both for estimating the global recombination rate in the painted lady and to investigate potential regional recombination rate variation and association with genomic features. The average, genome-wide recombination rate was 3.81 cM / Mb (W-chromosome excluded), but there was considerable inter-chromosomal variation (2.21 - 8.00 cM / Mb; Table S2, Figure S3), with a significantly higher rate on shorter chromosomes than on longer chromosomes (Spearman’s rank correlation, *ρ* = -0.83, p-value = 6.51×10^−07^;Figure 4). The recombination rate on the Z-chromosome was 3.09 cM / Mb, lower than the average unweighted autosomal rate. However, the recombination rate on the Z-chromosome was not lower than expected given the overall negative correlation between recombination rate and chromosome size (Figure 4A). Besides the negative association between chromosome size and recombination rate, we also found significant negative associations between chromosome size and GC-content (*ρ* = -0.65, p-value = 8.35×10^−05^) and repeat density (*ρ* = 0.77, p-value = 1.37×10^−06^), and a positive association with gene density (*ρ* = 0.68, p-value = 2.63×10^−05^)(Figure 4B-D).

**Figure 4.**
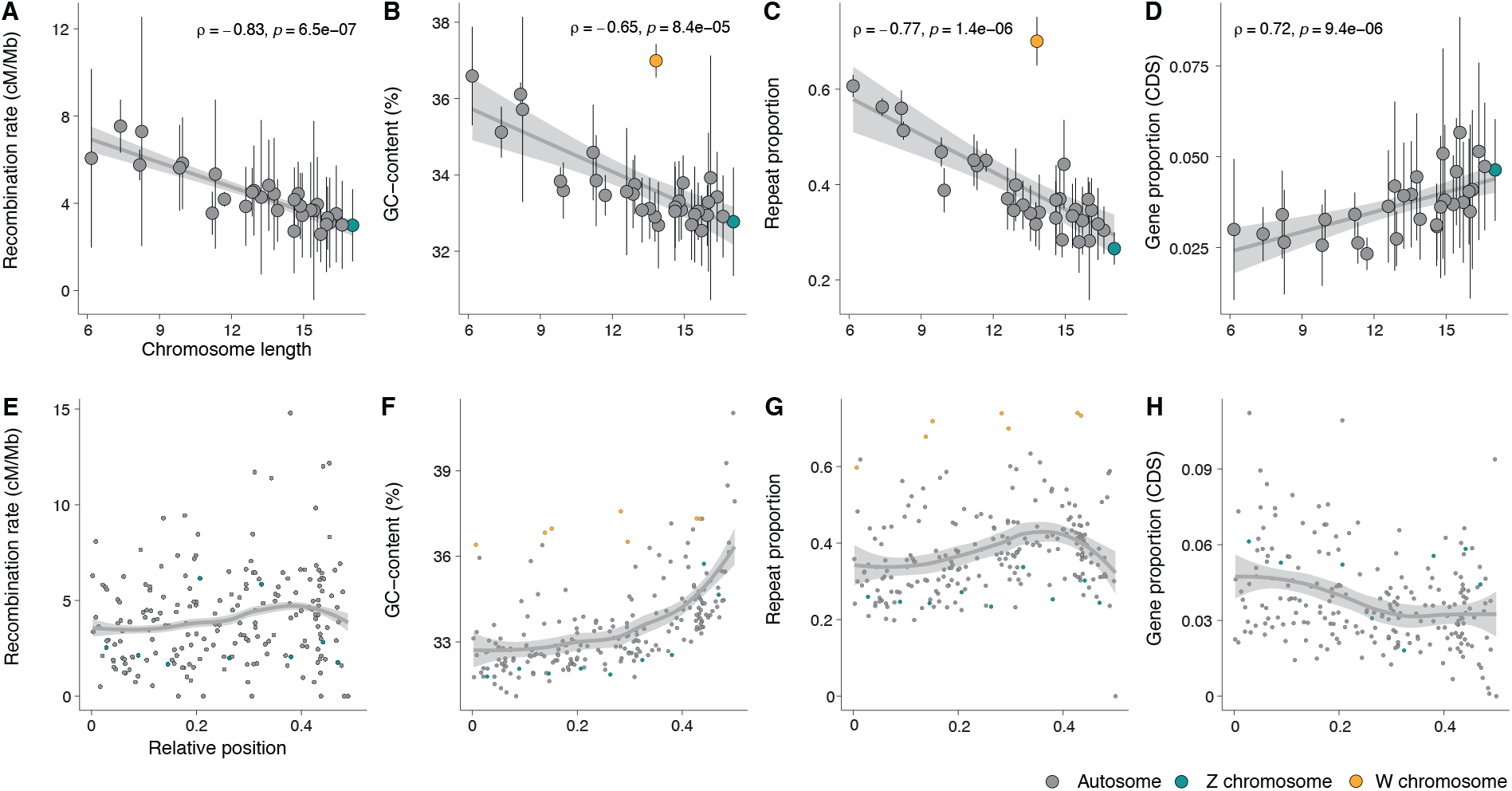
Top row: Associations between chromosome length and A) recombination rate, B) base composition, C) repeat and D) gene proportions. Chromosome length is given in megabases (Mb). Bottom row: Regional distribution of the recombination rate (E), base composition (F), repeat (G) and gene (H) density in 2 Mb windows along the chromosomes. All chromosomes were analyzed jointly and the x-axis shows the relative position (proportion of chromosomal length) from the center of the chromosomes.

#### Intra-chromosomal variation in recombination rate and associations with genomic elements

To quantify potential regional variation in recombination rate within chromosomes, we estimated the recombination rate in 2 Mb non-overlapping windows along each individual chromosome. The average rate across windows was similar to both the global rate estimate across chromosomes (4.05 +/- 2.45 cM / Mb) and the overall chromosome level estimates (2.58 - 7.53 cM / Mb, W-chromosome excluded). The recombination rate estimates for individual windows ranged between 0 - 14.79 cM / Mb (Figure S3, Table S2) and visual inspection revealed a bi-modal distribution with reduced recombination rate in the center of chromosomes and towards chromosome ends (Figure 4 E-H). To test this observation formally, we analyzed the difference in recombination rate between bins representing five relative distance intervals from the center of the chromosome for all chromosomes combined and found that the recombination rate was significantly lower in the center (first bin), significantly higher in the flanking terminal (fourth) regions and then again lower at the terminal end (Wilcoxon rank sum tests, p-values = 3.70×10^−02^ - 6.30×10^−15^; Figure S4).

To assess potential relationships between the recombination rate and genomic features in more detail, we first investigated different associations between the window-based recombination rate estimates and variation in nucleotide composition and proportions of different TEs and genes. The W-chromosome was excluded from this analysis since it is non-recombining in Lepidoptera. We found that the GC-content increased towards the ends of chromosomes and was positively associated to the regional recombination rate (*ρ* = 0.32, p-value = 3.68×10^−06^). Gene density was homogeneous across chromosomes, with only a minor increase towards the chromosome center, and was negatively associated with the recombination rate (*ρ* = - 0.19, p-value = 7.27×10^−03^). We found a significant positive association between the overall repeat proportion and the recombination rate (*ρ* = 0.35, p-value = 3.48×10^−07^; Figure 5), and this pattern was consistent for all repeat classes, but strongest for SINEs (*ρ* = 0.42, p-value = 2.63×10^−10^) and weakest for LTRs (*ρ* = 0.14, p-value = 4.04×10^−02^). The association between recombination rate and proportion of LTRs was, however, not significant when only including autosomes (*ρ* = 0.11, p-value = 11.04×10^−02^; Figure 5, Figure S5).

**Figure 5.**
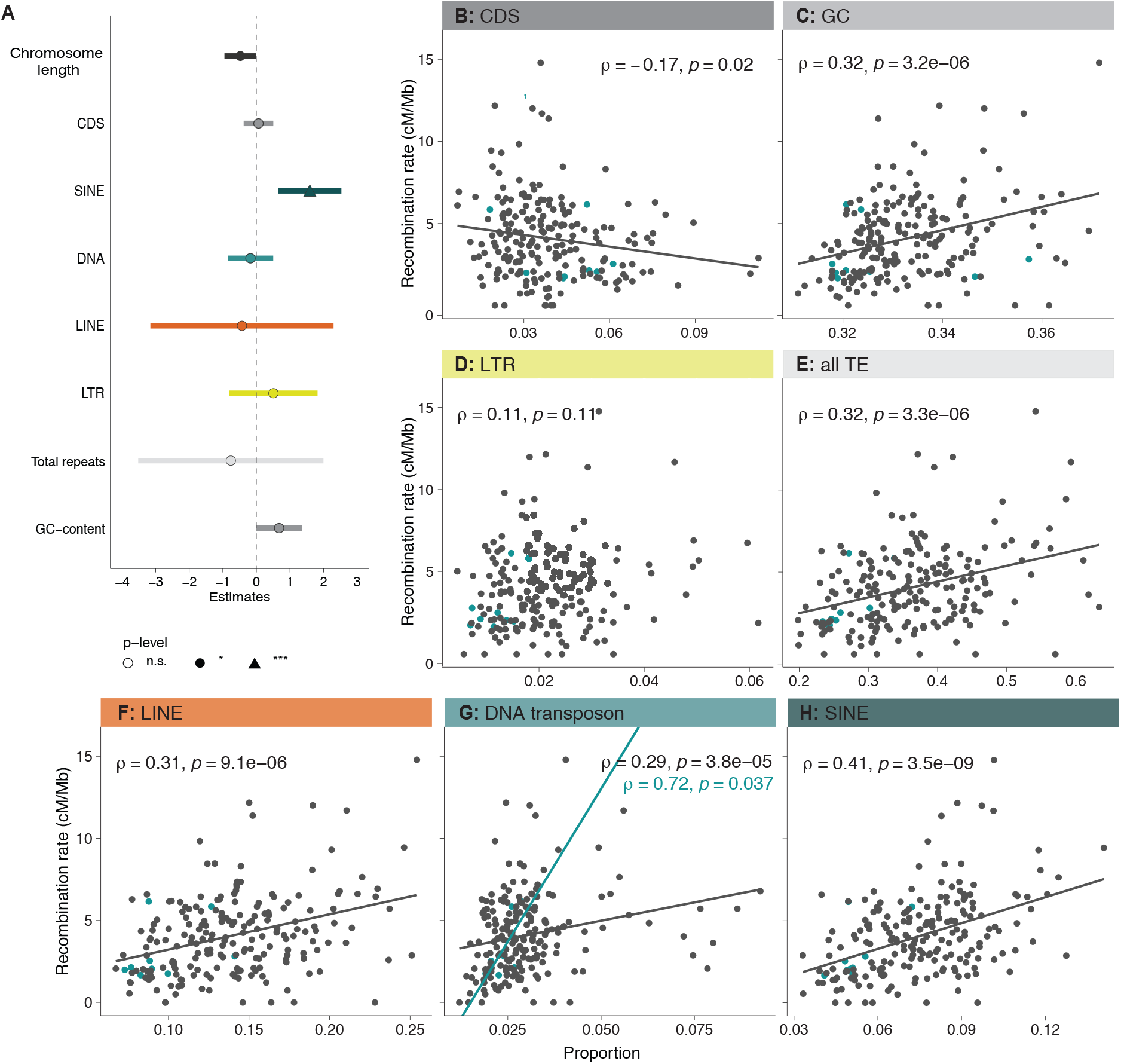
Correlation between recombination rate and density of genomic features. A) Summary of the linear model with regional recombination rate as the response variable. Each explanatory variable in the model is listed along the Y-axis and the relative estimated effect (X-axis), and error intervals are indicated with horizontal bars for each variable. B - H) Associations between the regional recombination rate and specific genomic features. Linear regression lines, Spearman’s correlation coefficients (*ρ*) and the corresponding *ρ*-values for significant analyzes are given. Gray dots/lines indicate autosomal regions and turquoise dots/lines Z-chromosome linked regions

To disentangle the relative strength of associations between the regional recombination rate and genomic features, a multiple linear model was implemented with recombination rate as the dependent variable. As explanatory variables we used chromosome length, chromosome type, GC-content, proportion of genes (CDS) and proportions of all different classes of TEs. We found that the regression model was significant (*df* = 197, *F* = 7.73, p-value = 5.36×10^−09^) and explanatory variables in the model accounted for 21% of the variation in recombination rate (*R*2 = 0.24, *adjR*2 = 0.21). Most of the variation was explained by the positive association with the proportion of SINEs (Estimate 1.59, p-value = 9.75×10^−04^) and the negative association with chromosome size (Estimate -0.48, p-value = 4.88×10^−02^; Figure 5, Table S3).

Finally, we explored whether gene expansions could be associated with other genomic features, and we therefore compared TE abundance in the regions with and without gene gains. The mean densities of LTRs, LINEs and DNA transposons were higher in regions with gene gains (Wilcoxon rank sum test, p-value 3.1×10^−03^ - 6.0×10^−04^; Figure S6), as was mean GC-content (p-value 3.0×10^−02^). The gene densities or recombination rates did not differ between regions with or without gene gains (Wilcoxon rank sum tests, p-values = 9.1×10^−01^ - 8.0×10^−01^; Figure S6).

## Discussion

### General

Here we present detailed results on the genomic architecture and regional recombination rate variation in the painted lady. The data paves the way for understanding the interplay between molecular mechanisms and micro-evolutionary processes shaping the genome of butterflies in general and provide the first insights into the links between genomic features and the unique lifestyle of this species. The rapid technological advances and dropping costs of DNA-sequencing methods have led to a staggering development rate of high-quality genome assemblies, including many butterfly species (Celorio-Mancera et al., 2021; Gu et al., 2019; Li et al., 2015; Smolander et al., 2022; Yang et al., 2020), and the availability of genomic resources will probably increase almost exponentially in the near future, as a result of the Darwin tree of Life (/https://www.darwintreeoflife.org/), the European Reference Genome Atlas (ERGA; https://www.erga-biodiversity.eu/) and other similar initiatives. However, detailed and curated genome annotation data are more time-consuming and expensive to generate and therefore still limiting comparative/population genomic and genotype-phenotype association approaches, not the least in butterflies (Davey et al., 2017; Hill et al., 2019; Van Belleghem et al., 2017). Another limiting factor for understanding both genome architecture in general, the relative effects of random and selective forces on sequence evolution and maintenance/loss of genetic diversity, and divergence processes, is that detailed recombination rate data are both laborious and time-intensive to gain, especially for natural populations. As a consequence, high-density recombination maps are still lacking for the vast majority of wild species where genome assemblies are now available. The detailed annotation information and the high-density linkage map for the painted lady developed here, therefore provide opportunities for both comparative studies on genome structure organization, population genomic- and micro-evolutionary investigations in the entire Lepidoptera clade.

Chromosome numbers have been shown to vary considerably between different butterfly and moth species; the haploid chromosome counts range from 5 to 223 (de Vos et al., 2020; Lukhtanov, 2015). In agreement with previous data (Zhang et al., 2021), both the linkage map and the DToL genome assembly clearly showed that the painted lady has a total haploid chromosome count of 31. We confirmed high levels of synteny and gene order collinearity between the painted lady and the silkmoth, and the lineage specific chromosome fusions characterized before in the postman butterfly (Davey et al., 2017). Hence, similar to other nymphalid butterflies, the painted lady has retained the inferred ancestral lepidopteran karyotype (Ahola et al., 2014). The annotation procedure revealed that the painted lady harbors a gene set (n = 13,161) close to the suggested core set in Lepidoptera (Challi et al., 2016; Li et al., 2019) and a relatively low overall TE content. However, the TE content was significantly higher and the gene density lower on smaller chromosomes. A clear outlier for gene density and TE content was the W-chromosome. While having a size equal to an average autosome, the W-chromosome demonstrated very specific features; both a significantly higher overall proportion of TEs, a larger fraction of longer TEs, and a different distribution of repeat classes compared to other chromosomes. Similar to the silkmoth and julia heliconian (*Dryas iulia*), the W-chromosome in the painted lady had a significantly higher proportion of LTRs and LINEs (Lewis et al., 2021; Mita et al., 2004). The proportion of SINEs was however much smaller on the W-chromosome than on the autosomes and the Z-chromosome. A lack of protein coding genes, like we observed on the painted lady W-chromosome, has also been observed in the silkmoth (Abe et al., 2008; Mita et al., 2004), and is likely a consequence of the general degradation process of the non-recombining sex-chromosome (Bachtrog, 2013). The higher accumulation of TEs is also an expected consequence of recombination suppression and comparatively low effective population size (*N*_*e*_) of the W-chromosome (1*/*4 of the autosomes at equal sex-ratios), both as a consequence of Müllers ratchet and since the overall efficiency of selection against TE insertion is reduced for non-recombining chromosomes (Bachtrog, 2013). The Z-chromosome is generally highly conserved in Lepidoptera (Fraïsse et al., 2017) and it is the largest of all the painted lady’s chromosomes. We did not find any significant differences in gene or TE content on the Z-chromosome compared to the autosomes.

### Gene family analysis

Gene family expansions can provide the raw material for both neo- and sub-functionalizing evolutionary directions, and the rate of gene duplication can be significantly higher than the rate of function-altering single nucleotide mutations (Lipinski et al., 2011). However, most gene duplication events are probably deleterious (Loehlin and Carroll, 2016) or effectively neutral, leading to a low probability of fixation of novel gene copies (Emerson et al., 2008). We found a comparatively low proportion of lineage-specific gene duplications in the painted lady, which could be a consequence of the large *N*_*e*_ of the species (Garcìa-Berro et al., in prep), which translates to efficient selection against slightly deleterious variants. The majority of the (comparatively few) significant gene expansions in the painted lady lineage clustered on single chromosomes - only a single gene family had expanded and dispersed across multiple chromosomes - suggesting that unequal crossing over has been the main mechanism behind gene family expansions.

The painted lady has extraordinary life-history characteristics and has become a quickly uprising complementary model organism for studying insect migration. Over most of the almost cosmopolitan distribution range (Shields, 1992), the painted lady individuals annually complete a multigenerational migratory circuit that can span many thousand kilometers in total, and single individuals can migrate > 4, 000 kilometers during lifetime (Talavera and Vila, n.d.). In addition, the painted lady is a polyphagous generalist that utilizes > 300 different larval host-plants in 11 plant families (Celorio-Mancera et al., n.d.; Nylin et al., 2014) and, in contrast to other migratory butterflies like the monarch and the red admiral (*Vanessa atalanta*), the painted lady is non-diapausing (Shields, 1992). This unique combination of life-history traits is accompanied by high levels of heterozygosity and presumably a very large effective population size (Garcìa-Berro et al., in prep). The genetic underpinnings of migratory behavior have only been preliminarily characterized for a handful of insect species (Kang et al., 2004; Zhu et al., 2009) and have not been studied in painted lady before. The dissection of potential associations between genetic (and epigenetic) variants and complex phenotypes like ‘migratory behavior’ obviously requires a combination of multiple approaches. As the first step to understanding lineage-specific characteristics of the painted lady, we here focused on gene family evolution. Our results showed a limited number of genes with significant copy number expansions unique to the painted lady lineage. The expanded gene families were mainly associated with functions related to the transport of fatty acids, protein metabolism, and muscle structure and activity. Since migratory insects mainly use fat as an energy resource during migration (Landys et al., 2005; Murata and Tojo, 2013; Srygley and Dudley, 2008; Weber, 2009), both the capacity to build up fat deposits and efficient sequestration of fatty acids have likely been under strong selection in the painted lady. Likewise, enhanced muscle structure and function should be advantageous for long-distance migrants compared to sedentary species.

Therefore, efficient fine-tuning and optimization of fatty acid metabolism and increased muscle sustainability during migration could have been aided by the expansion of specific gene sets involved in those processes.

Besides the obvious advantages of having efficient energy metabolism and high-functioning flight machinery, long-range migrants will also benefit from utilizing a multitude of different host plants since they will encounter dramatically different habitats, both during the lifespan of single migratory individuals and between consecutive generations. In contrast to the monophagous monarch butterfly, the painted lady can utilize a wide range of host plant species (Ackery, 1988; Nylin et al., 2014), an adaptation that probably has been coupled to strong selection on genes involved in detoxification of secondary metabolites. We found that two of the significantly expanded gene families in the painted lady (UDP-glycosyltransferase, carboxylesterase) were associated with detoxification and polyphagy (Breeschoten et al., 2022; Hatfield et al., 2016; Nagare et al., 2021). The UDP-glycosyltransferase superfamily includes Lepidoptera-specific subfamilies associated with a variety of functions, such as affinity for plant secondary metabolites (Huang et al., 2008; Luque et al., 2002). In the painted lady larvae, one UDP-subfamily is upregulated in response to utilization of an extended range of host-plants (Celorio-Mancera et al., n.d.). Copy-number expansions of these detoxifying gene families could have allowed the painted lady to increase the range of host plants that can be utilized and consequently paved the way for developing the non-diapausing, multigenerational, long-distance migratory lifestyle. The wide range of habitats that long-distance migratory species encounter also probably means that they are exposed to many more different pathogens than sedentary species. Our analysis revealed that the

Lepidoptera-specific gene *moricin*, associated with inducible antimicrobial peptides (Hara and Yamakawa, 1995), was significantly expanded in the painted lady. An increase in the number of *moricin* copies could have increased the efficiency of defense against a larger suite of pathogens.

The genetic basis of migratory behavior has been investigated in some detail in the monarch butterfly. A combination of approaches has identified candidate genes associated with orientation, chemoreception and regulation of the circadian clock (Zhan et al., 2014; Zhu et al., 2009). Migratory behavior has evolved independently multiple times within the Papilionoidea clade (Chowdhury et al., 2021) and in the *Vanessa* genus (Wahlberg and Rubinoff, 2011), and the life histories of the monarch butterfly and the painted lady are distinct. However, long-distance migration should put selective pressure on similar traits (e.g. navigation, energy metabolism, muscle endurance), and it is therefore possible that specific gene categories have been under selection in independent lineages. We therefore expanded the analysis to include gene-family expansions that were shared between the painted lady and the monarch in our sample set. Significantly expanded gene families were enriched for functions associated with various metabolic processes, defense against pathogens and neuronal activity, all of which are straightforward to intuitively associate with migratory behavior. One gene family with expanded copy numbers in both species and an especially pronounced increase in the painted lady was a family of vacuolar ATPases. The ATPases are ATP-dependent proton pumps involved in membrane transports, and they have been shown to affect for example ion transport in insects (Wieczorek et al., 2009). Given the unique expansion of this gene family in both species, we speculate that copy number increase could be involved in flight muscle coordination and/or ion transport for maintenance of homeostasis during long periods of flight.

In this study, we get a first glimpse of the specific genes that have undergone copy number expansions in painted lady specifically and independently in the two migratory species. The functions associated with the expanded gene families can be coupled to the evolution of long-distance migratory behavior. However, further studies on larger species sets with independent migratory and sedentary sister species pairs, in combination with detailed intraspecific population genetic analysis and functional verification experiments will be necessary to dissect the genetic underpinnings of migratory behavior in butterflies in detail.

### Patterns of recombination rate variation

Detailed data on recombination rate variation are crucial for understanding the relative effects of random genetic drift and selection on levels of genetic diversity and disentangling the evolutionary forces shaping genetic divergence between incipient species. Understanding how recombination breaks down linkage disequilibrium between physically linked regions is also important for the efficient design of association studies aimed at coupling genetic variation to phenotypic traits. Despite these important contributions to evolutionary genomics research, detailed recombination maps are only available for a handful of butterfly species (Beldade et al., 2009; Celorio-Mancera et al., 2021; Davey et al., 2017; Rosser et al., 2022; Smolander et al., 2022; Tunström et al., 2021). In some cases, linkage maps have been used to improve and/or verify the correctness of physical genome assemblies, but analysis of the recombination rate has not been thoroughly assessed in many butterfly species. Here we developed a high-density linkage map based on segregation information in a pedigree with 95 offspring. The map contained > 1, 300 ordered markers and the overall density was > 3 markers per Mb. Despite being based on a single pedigree, the genetic map developed here revealed a recombination landscape in strong agreement with what has been observed in other butterflies (Davey et al., 2017; Martin et al., 2019). This indicates that the painted lady genetic map reflects the historical recombination landscape in the species well.

We estimated the genome-wide average recombination rate in the painted lady to be 3.81 - 4.05 cM / Mb, dependent on the method applied. The global rate was in the lower end of recombination rate estimates from other Lepidoptera species, which have been in the range from 2.97 - 4.0 cM / Mb in the silkmoth (Yamamoto et al., 2008; Yasukochi, 1998) to 5.5 - 6.0 cM / Mb in different *Heliconius* species (Jiggins et al., 2005; Tobler et al., 2005). We found a significant negative association between chromosome length and the recombination rate in the painted lady. This is a consistent pattern found across many organism groups and likely a consequence of that at least one crossover event is necessary for correct segregation of chromosomes during meiotic division in the recombining sex, leading to a higher recombination rate per unit length for shorter chromosomes (Haenel et al., 2018; Kawakami et al., 2017; Martin et al., 2019). Butterflies and moths are holocentric, i.e. they lack distinct centromere regions which means that the spindle fibers can attach ‘anywhere’ along the chromosomes during cell division. This might lead to an expectation of a more uniform distribution of recombination events along chromosomes in holocentric species if crossovers occur randomly. The window-based analysis in the painted lady revealed a bimodal distribution of recombination events along chromosomes, with a significantly higher rate in regions close to, but not directly at, chromosome ends. This distribution is in agreement with previous observations, both in Lepidoptera and in other animals with different centromere types (Haenel et al., 2018; Martin et al., 2019). A possible explanation for this pattern could be mechanical or tension interference between chiasmata when > 1 recombination event occurs on the same chromosome during the same meiotic division (Haenel et al., 2018). However, in the holocentric *Caenorhabditis elegans*, the number of recombination events is limited to precisely one per chromosome per meiosis, but there is still a strong bimodal pattern of recombination rate variation along chromosomes in this species (Barnes et al., 1995). An alternative explanation for the bimodal distribution of recombination events along chromosomes could be that synaptonemal complexes are directed towards specific physical positions when the telomeres attach to the nuclear wall (Scherthan et al., 1996). As indicated above, we also found that the recombination rate dropped significantly at the far ends of the chromosomes in the painted lady. This reduced recombination rate at chromosome ends is also consistent with earlier observations and could potentially be attributed to selection against synaptonemal complex formation at chromosome ends, due to a higher risk of ectopic recombination in these generally repeat-rich regions (Smith and Nambiar, 2020).

Since recombination is directly associated with the efficacy of selection, a negative correlation between the regional recombination rate and a number of repeats would be expected if TE insertions predominantly are deleterious. Such associations have been observed in many organisms, although the relationship between TE-abundance and the recombination rate varies to some extent across species and different TE-classes (Kent et al., 2017; Rizzon et al., 2002). In the painted lady, we observed a significant positive association between TE-abundance and the regional recombination rate, predominantly driven by a strong effect of SINE density. An explanation for the strong association between SINE density and recombination rate could be SINE-mediated recombination, as has for example been described in humans (Deininger and Batzer, 1999), but we can not exclude other factors affecting both recombination rate and the proliferation efficiency of SINEs. For example, both synaptonemal complexes and SINE insertions might be directed towards regions of more open chromatin structure. One interesting observation was the radically different distribution of TE-classes on the W-chromosome in the painted lady, with a very low frequency of SINEs as compared to the autosomes and the Z-chromosome. Female heterogamety has been conserved across both Lepidoptera and Trichoptera and the lepidopteran W-chromosome probably developed as a result of an ancestral Z-chromosome to autosome fusion > 90 million years ago (Fraïsse et al., 2017). The lack of functional protein coding genes and the significant enrichment of specific TEs suggest that the W-chromosome has been non-recombining over most of that time span. The absence of SINEs on the W-chromosome, and the strong positive association between SINE density and recombination rate on the autosomes and the Z-chromosome, hence suggests that SINEs might be able to hijack the recombination machinery and mediate their own proliferation via double-strand breaks.

In contrast with results from similar studies in other organism groups (Apuli et al., 2020; Kawakami et al., 2014), we observed a negative association between the recombination rate and gene density. This is likely a consequence of the strong association between recombination rate and chromosome size, since the association with gene density was insignificant when chromosome size was included as an explanatory variable. The observed weak positive association between GC-content and recombination is in agreement with the limited effect of GC-biased gene conversion (gBGC) in butterflies (Boman et al., 2021). We did not find any association between recombination rate and the presence of extended orthogroups, which would be expected if gene duplication is associated with unequal crossing-over. This could possibly be a consequence of the more efficient removal of deleterious duplications in regions with higher recombination rate. However, repetitive elements can trigger ectopic recombination which can explain the observed significant positive association between gene gains and density of LTRs, LINEs and DNA elements in the painted lady.

## Conclusions

In this study, we present detailed annotation and recombination rate information for the painted lady butterfly (*Vanessa cardui*), a species with a remarkable life-history traits such as long distance migration, continuous direct development and a capacity to utilize many different types of larval host plants. We analyzed lineage-specific gene family expansions and found that expanded genes were mainly associated with fat and protein metabolism, detoxification and defense against pathogens. A detailed TE-annotation revealed that several TE-classes were positively associated with the presence of gained genes, potentially indicating their involvement in ectopic recombination. Recombination rate variation was negatively associated with chromosome size and positively associated with the proportion of short interspersed elements (SINEs). We conclude that the genome structure of the painted lady has been shaped by a complex interplay between recombination, gene duplications and repeat activity and provide the first set of candidate genes potentially involved in the evolution of migratory behavior in this almost cosmopolitan butterfly species.

## Methods

### Linkage map

#### Sampling and DNA-extraction

Offspring from one painted lady female were reared on thistles (*Cirsium vulgare*) in the greenhouse until pupation. The bursa copulatrix of a female was examined and only one spermatophore was detected, indicating that a single male had sired all offspring. The offspring were snap frozen in liquid nitrogen and stored in −20°C until DNA extraction. DNA was extracted from thorax tissue of the female and an abdominal segment of the offspring pupae, using a modified high salt extraction method (Aljanabi, 1997). The quality of the DNA was analyzed with Nanodrop (ThermoFischer Scientific) and the yield was quantified with Qubit (ThermoFischer Scientific). Extracted DNA was digested with the restriction enzyme EcoR1 according to the manufacturer’s protocol, using 16 hours digestion time (ThermoFischer Scientific). DNA fragmentation was verified with standard gel electrophoresis. Digested DNA from 95 offspring with the highest yield and the dam was shipped to the National Genomics Infrastructure (NGI, see acknowledgements) in Stockholm for library preparation (standard protocol), individual barcoding and multiplex sequencing using 2 × 151 bp paired-end reads on one NovaSeq6000 S4 lane.

#### Building the linkage map

The quality of the raw reads was assessed with FastQC (Andrews et al., 2012). The reads were filtered using the Stacks2 modules `clone_filter` to remove PCR-duplicates and `process_radtags` to filter for quality. We evaluated phredscore in sliding windows covering 15% of the read length and removed reads with mean score below 10 (Catchen et al., 2013). Removal of reads with unassigned bases and truncation to 125 bp was done using option `-c`, and `--disable_rad_chec` was applied to keep reads with incomplete RAD-tags.

We mapped the filtered reads to the previously published genome assembly (Lohse et al., 2021) using the bwa mem algorithm (Li, 2013) with default options. Resulting bam files were sorted with samtools sort (Li et al., 2009) and filtered with `samtools view –q 10` (only reads with mapping quality score above 10 were retained). A custom script was applied to retain reads with unique hits only. The mapping coverage was analyzed with Qualimap (Okonechnikov et al., 2015). The offspring were defined as females if the coverage on the Z-chromosome was *<* 75% of the average coverage over all chromosomes and as males if the coverage was > 75%. Samtools mpileup was used for variant calling using minimum mapping quality (`-q`) 10 and minimum base quality (`-Q`) 10 (Li et al., 2009). The variants were then converted to likelihoods with Pileup2Likelihoods in LepMap3 using default settings (Rastas, 2017). The LepMap3 protocol (Rastas, 2017) with some modifications was used to construct the linkage map (Supplementary methods 1).

### Genome annotation and whole genome statistics

#### Genome assembly statistics

With very few exceptions, the order of markers in the linkage map was in agreement with the physical order in the assembly. We therefore did not make any corrections to the physical assembly before further analysis. Standard genome assembly summary statistics were calculated for the genome assembly after linkage map verification, using the QUAST suite (Gurevich et al., 2013). For the subsequent analysis we excluded unassembled haplotigs from the genome assembly and retained all the other scaffolds.

We used MCScanX (Wang et al., 2012) to detect syntenic blocks between the painted lady genome assembly on the one hand and the silkmoth and the the postman butterfly on the other. We downloaded the annotation for the silkmoth assembly from SilkBase (/https://silkdb.bioinfotoolkits.net/) and the 2.5-version of the postman annotation from LepBase (download.lepbase.org/v4/, downloaded 2021-06-21). BLAST was used for primary alignment and we used a custom script to select the five hits with the highest E-values and used them as input for the MCScanX. The CIRCOS library (Krzywinski et al., 2009) was used for visualization of the results.

#### Gene and repeat annotation

The annotation of the painted lady genome assembly was performed using MAKER version 3.00.0 (Holt and Yandell, 2011) iteratively in three steps. In the first step, we mapped previously available transcriptomic evidence data from the painted lady based on wing tissue (Connahs et al., 2016)(accessed on 2020-05-15) and masked all known repeats. RepeatMasker version 4.0.3 (Smit et al., 2015) was used within the MAKER pipeline with a manually curated Lepidoptera repeat library (Talla et al., 2017) serving as a reference. The first MAKER round produced a set of gene models, which were quality controlled using Annotation Edit Distance (AED) statistics. AED quantifies congruency between a gene annotation and its supporting evidence. We discarded gene models with AED scores higher than 0.5 (50% of the gene model length not matching the corresponding evidence sequence) using custom scripts. The retained gene models were provided as a training set for the second run of MAKER.

The second iteration of MAKER was run to generate gene models using the *ab initio* gene predicting algorithm implemented in SNAP (Korf, 2004). For the last step in the MAKER pipeline, gene models predicted by SNAP and additional protein evidence from the Uniprot database (https://www.uniprot.org/; accessed 2021-04-01) were used. A set of Lepidoptera proteins from the Swiss-prot section of the Uniprot database were downloaded and manually curated. All genes from the curated set were included while only fully sequenced nuclear proteins with predicted functions from the non-curated gene set were included (custom scripts were used for selection). This selection resulted in 36,907 proteins. Finally, all obtained evidence and *ab initio* predicted genes were merged resulting in 18,860 gene models. Resulting genes were renamed using MAKER supplementary scripts.

### Additional curation

The gene models constructed by MAKER were filtered based on standard options; discarding gene models with AED and eAED scores *<* 0.5 and/or length *<* 50 amino acids. To search for functional domains in putative genes, we used InterProScan with default settings (Jones et al., 2014). A number of TE-related domains were detected within gene models indicating a need for a more detailed transposable element annotation. The repeat library was extended by adding evidence from the current RepBase for all Arthoropoda (Bao et al., 2015) and curated repeats from the monarch butterfly (Zhan and Reppert, 2013). RepeatModeler and RepeatMasker (Holt and Yandell, 2011) were thereafter run again with the updated repeat database.

Coordinates of newly identified repeats were intersected with gene model positions using BEDTools (Quinlan and Hall, 2010) and we removed gene models that overlapped more than 50% of the length of a repeat. We then searched for keywords in the InterProScan domain output and removed genes containing at least one TE domain. For the W-chromosome, we manually curated the InterProScan output and found no functional information for any genes. The filtering resulted in a total set of 13,161 genes, including 12,209 genes with preliminary assignments in eggnog (Huerta-Cepas et al., 2019). The entire painted lady gene set represented 89.9% of the complete BUSCO arthropod gene set (Manni et al., 2021).

### Gene family evolution

We investigated gene family evolution in the painted lady by comparing our obtained gene annotations with other annotated nymphalid genomes available on Lepbase. Protein fasta files for eight other nymphalid species - squinting bush brown (*Bicyclus anynana*), monarch (*Danaus plexippus*), postman (*Heliconius melpomene melpomene*), red postman (*Heliconius erato lativitta*), common buckeye (*Junonia coenia*), ringlet (*Maniola hyperantus*), speckled wood (*Pararge aegeria*) and Kamehameha butterfly *(Vanessa tameamea)* - were downloaded from Lepbase (http://download.lepbase.org/v4/sequence/: accessed 2021-06-21; Supplementary Table S1). The annotated gene sets in each species were filtered to only include one transcript per gene. To cluster the annotated genes into orthogroups and infer species specific orthogroups and gene duplications, OrthoFinder/2.5.2 (Emms and Kelly, 2019) was used with default settings. The total gene counts for each orthogroup and species from OrthoFinder was used as input to estimate gene family expansions and contractions with the software BadiRate, using the maximum likelihood option and the birth/death/innovation (BDI) model (Librado et al., 2012). We used the species tree obtained with OrthoFinder as input, with additional conversions using Tree as implemented in ete3 (Huerta-Cepas et al., 2016).

For each orthogroup identified in OrthoFinder, different models reflecting the evolution of the gene families were tested. The null model (Global rate model) assumes a uniform rate of gene gains/losses for all branches in the provided species tree. Alternative models were specified as follows; i) to detect gene family changes specific to the painted lady, a distinct branch rate was specified in the painted lady with all the other branches evolving at uniform, background rates, and, ii) the terminal branches of the two distinct migratory species, the painted lady and the monarch, were set at a common rate, differing from all other taxa in the data set. The rationale behind the last setting was to allow for identification of gene family expansions shared between the painted lady and monarch butterfly. Each model was run twice and the replicate with highest likelihood for each model was used for model comparisons.

Likelihoods of all models were compared using Aikaike’s Information Criterion (AIC) (Akaike, 1974), calculated as 2*K −* 2 log *L*, where *K* is the number of parameters and log *L* is the logarithm of the likelihood of the model. The orthogroups where the alternative models in BadiRate inferred gene gains > 0 with the lowest AIC, were used for analysis of functional enrichment. BadiRate was partly run with a modified version of the R-package BadiRateR (BadiRate) and custom scripts. We visualized the genomic location of genes belonging to extended orthogroups identified in BadiRate with custom bash scripts and PhenoGram (Wolfe et al., 2013), using gene names and positions from the annotation gff file.

#### Gene ontology enrichment

Potential enrichment of functional categories in the significantly expanded gene sets was analyzed using the Bioconductor package topGO version 2.44.0 (Alexa and Rahnenfuhrer, 2021) in R version 4.1.0 (R Core Team (2021), 2021). A custom database was generated based on the annotated gene set with gene ontology (GO) terms associated to the categories biological process, cellular component and molecular function. Since the gene set of interest is based on gene counts, the enrichment test was performed with Fisher’s exact test using the default algorithm (”`weight01`”) which accounts for the hierarchical structure of the GO-terms (Alexa et al., 2006). This means that the resulting tests were not completely independent and correcting for multiple testing might be over-conservative. We still adjusted the p-values with Benjamini-Hochberg’s method of multiple test correction (Benjamini and Hochberg, 1995).

### Recombination rate analysis

#### Chromosome level analysis

Global and chromosome-specific recombination rates were estimated by dividing the linkage map length (unit = cM) with the physical length (bp) of the corresponding part (whole genome or individual chromosome) of the physical genome assembly. Regional recombination rates were estimated with local linear regression in 2 Mb non-overlapping windows containing 2 or more markers using the R-package MareyMap (Rezvoy et al., 2007).

#### Window-based analysis

We quantified the spatial distribution of various genomic features along the painted lady genome using custom scripts (available on GitHub, see Data Accessibility). Positions of the specific TE classes were accessed from the RepeatMaker output file and positions of genes were taken from the annotation file from MAKER. For each 2 Mb window, we calculated the density (fraction of window covered by element / window length) of genes, TEs in total, LTRs, SINEs, LINEs and DNA-transposons with in-house developed scripts. For the genes belonging to extended orthogroups, we integrated the list of the extended orthogroups from BadiRate, gene names from OrthoFinder and positions from MAKER. All 2 Mb windows of the genome were assigned to bins; one bin containing windows without gene gains and the other with windows containing at least one gained gene. Potential differences in densities of genomic features between bins were assessed with two-sided Wilcoxon rank sum tests in R (Bauer, 1972).

To characterize the associations between recombination rate and specific genomic elements, correlation tests were performed with the cor.test function in R using Spearman’s rank correlation (Best and Roberts, 1975), after testing for deviation from normal distribution with the Shapiro-Wilk normality test (Royston, 1982). We then applied the `lm` function in R to explore the relationships between the recombination rate as response variable and chromosome length, relative position on the chromosome, density of genes, density of different repeat classes, and GC-content as explanatory variables. Prior to the latter analysis, explanatory variables were scaled and centered, by subtracting the mean and dividing by the standard deviation of the variable. We used the R-package `ggplot2` for visualizations (Wickham, 2009).

## Data access

All raw data have been submitted to the European Nucleotide Archive (ENA accession number XXXX). Scripts are available in the GitHub repository (GitHub LINK XXXXX).

## Acknowledgements

Financial support for this project was provided by FORMAS (Research grant 2019-00670 to N.B.) and The Swedish Collegium for Advanced Science (Natural Sciences Programme, Knut and Alice Wallenberg Foundation, Postdoc funding for D.S.). R.V. was supported by the grant PID2019-107078GB-I00 funded by MCIN/AEI/10.13039/501100011033. G.T. was supported by the grant PID2020-117739GA-I00 funded by MCIN/AEI/10.13039/501100011033 and by “La Caixa” Foundation (ID 100010434) through the grant LCF/BQ/PR19/11700004. The authors acknowledge support from the National Genomics Infrastructure in Stockholm funded by Science for Life Laboratory, the Knut and Alice Wallenberg Foundation and the Swedish Research Council, and SNIC/Uppsala Multidisciplinary Center for Advanced Computational Science for assistance with massively parallel sequencing and access to the UPPMAX computational infrastructure.

